# Knowledge Discovery in Datasets of Proteomics by Systems Modeling in Translational Research on Pancreatic Cancer

**DOI:** 10.1101/2025.02.23.639474

**Authors:** Mathilde Resell, Elisabeth Pimpisa Graarud, Hanne-Line Rabben, Animesh Sharma, Lars Hagen, Linh Hoang, Nan T. Skogaker, Anne Aarvik, Magnus K. Svensson, Manoj Amrutkar, Caroline S. Verbeke, Surinder K. Batra, Gunnar Qvigstad, Timothy C. Wang, Anil Rustgi, Duan Chen, Chun-Mei Zhao

## Abstract

Knowledge discovery in databases (KDD) can contribute to translational research, also known as translational medicine, by bridging the gap between *in vitro* and *in vivo* studies and clinical applications. Here, we propose a ‘systems modeling’ workflow for KDD. This framework includes data collection of composition model (various research models) and processing model (proteomics) and analytical model (bioinformatics, artificial intelligence/machine leaning and pattern evaluation), knowledge presentation, and feedback loops for hypothesis generation and validation. We applied this workflow to study pancreatic ductal adenocarcinoma (PDAC). Through this approach, we identified the common proteins between human PDAC and various research models *in vitro* (cells, spheroids and organoids) and *in vivo* (mouse mice). Accordingly, we hypothesized potential translational targets on hub proteins and the related signaling pathways, PDAC specific proteins and signature pathways, and high topological proteins. Thus, we suggest that this systems modeling workflow can be a valuable method for KDD, facilitating knowledge discovery in translational targets in general and in particular to PADA in this case.

## Introduction

In translational research, various experimental models have been developed with advantages as well as limitations. In the field of human cancer modeling, *in vitro* models (e.g., cell lines and organoids) and *in vivo* models (e.g., mice) are widely used either alone or in combination. However, it should be noted that the cell lines used in 2D *in vitro* and spheroids in 3D *in vitro* experiments often diverge from their parental origins due to ongoing genetic modifications over the passages and adaptions to culture environments and that the tumor organoid models in 3D *in vitro* experiments can change in the composition of cell types over time. The mouse models, as *in vivo* experiments, have species-specific differences which are often questioned about accurately recapitulation of *de novo* human tumor development. Recently, we presented a data collection on pancreatic cancer by gathering and analyzing accurate data from the *in vitro* and *in vivo* models for translation research^1^. In this work, we further present a workflow of “systems modeling” as a process of developing abstract models of a system, with each model presenting a different view or perspective of that system, for knowledge discovery in databases (KDD). The workflow consists of a data processing model (i.e., proteomics), composition model (different experimental models), data mining models (data selection and integration, and functional enrichment analysis and artificial intelligence/AI/ML, AI/ML), and knowledge presentation model. We validated this workflow with particularly focusing on pancreatic cancer.

Pancreatic cancer refers here to pancreatic ductal adenocarcinoma (PDAC) which is the 3^rd^ deadliest cancer in developed countries. It is characterized by an insidious clinical syndrome with aggressive malignant and dismal outcomes with a 5-year survival rate of ≈10%^2,3^. Despite substantial research efforts, investment in PDAC has yet to yield significant improvements in both survival rates and quality of life (QoL) over the past decades. In clinical practice, PDAC is seldom diagnosed at a stage when curative surgery is viable, primarily due to the absence of early detection biomarkers, even among high-risk populations. A common treatment in patients with inoperable PDAC is chemotherapy which may offer modest benefits^4,5^. Current “experimental” treatments include immunotherapies and various targeted therapies, whereas palliative care focuses on symptom management to improve QoL ^6–9^.

The development of molecular targeted treatments for PDAC relies on a profound understanding of its molecular biology. Currently, identified genetic mutations that serve as potential genomic biomarkers for PDAC include the activation of the oncogene KRAS, the inactivation of tumor suppressor genes, the inactivation of genome maintenance genes, and alterations in genes specifically involved in the homologous recombination repair pathway ^10^. These mutations present both opportunities and challenges for targeted therapy. For example, the FDA approval of sotorasib for targeting the KRASG12C mutation has been applied to only 1-2% of PDAC patients who exhibited this specific mutation ^11^. Several molecules that demonstrated promising preclinical efficacy failed in clinical trials, likely due to off-target effects such as intolerability, adverse reactions and failure to achieve objective responses. Together, these represent the challenge of the ‘translational gap’ in the research on PDAC^12^.

## Results

### Workflow of systems modeling for KDD

We hypothesized that the translational gap in PDAC research arises from mismatches between research models and actual patient conditions. To address this, we aimed to develop and validate the systems modeling workflow designed to identify ‘translatable target’ for PDAC. Our workflow incorporated a comprehensive range of PDAC models, including murine PDAC cells, murine normal pancreas tissue, murine PDAC spheroids, murine PDAC organoids, murine pancreatic exocrine organoids, human PDAC organoids and human PDAC tumor tissue for data collection, particularly on proteomics. Analytic methods included bioinformatics and AI/ML. With filters in each step, the knowledge on potential translational targets was discovered, which led to new hypothesis generation, test and validation (Fig.1).

**Fig. 1.**
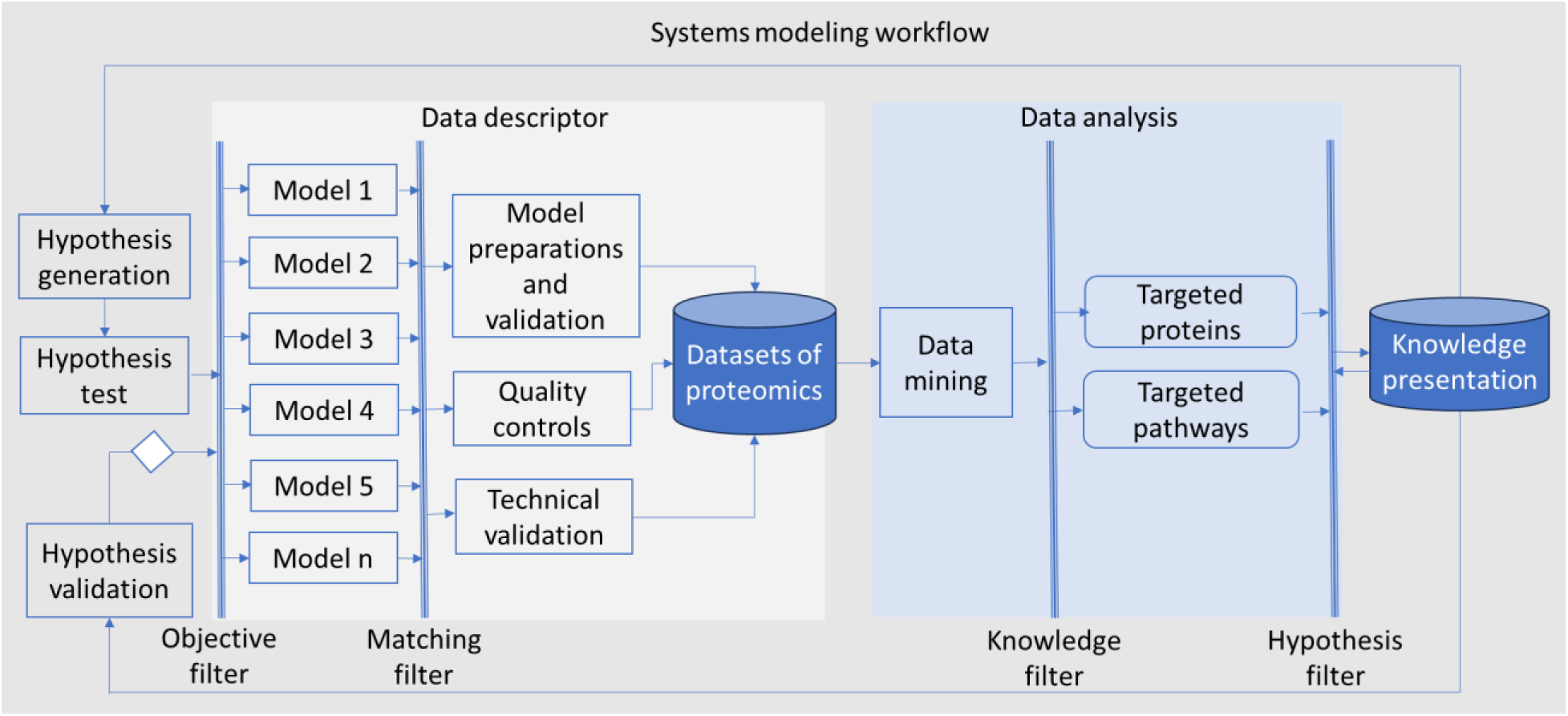
Schematic diagram of systems modeling. Hypotheses in translational medicine are formulated based on clinical observations (‘bedside’) and research findings (‘bench’). These hypotheses are subjected to ‘objective filter’ and then tested using one or more research models, clinical data and trials. In this study, five distinct models were employed, including murine PDAC cells, spheroids, organoids, orthotopic model, and human PDAC organoids. To address the translational gap, patient-derived materials were utilized as a ‘matching filter,’ ensuring that the models closely align with the human disease. The datasets included in this work comprised proteomic profiles from both research models and patient samples (showed in grey scare)^1^. The data analysis in this work (showed in light-blue square) included data mining with functional enrichment analysis and machine leaning. A ‘knowledge filter’, as a ‘human-in-the-loop’ element, was employed for interpreting data, and a ‘hypothesis filter’ was used for the knowledge discovery, e.g., targeted proteins and signaling pathways. The knowledge presentation in this work can lead to validation of hypothesis and generation of new hypotheses. This workflow is adaptable, allowing for the incorporation of other models and filters according to hypothesis, such as human cell lines, genetically engineered mouse models, tumor microenvironment components like fibroblasts and cancer-associated fibroblast organoids, and blood samples according to hypotheses.

### Knowledge presentation of common proteins across various research models and human PDAC

Common proteins were defined as the proteins presented in the intersection of all models, as visualized in the Venn diagram. Only proteins that passed the initial filtering within each model and were subsequently found in all six models were considered the common proteins in the study. The research models shared proteins with human PDAC in a range between ∼50-60% (Fig. 2), highlighting both the existence of a translational gap and the potential for successful translation.

**Fig. 2.**
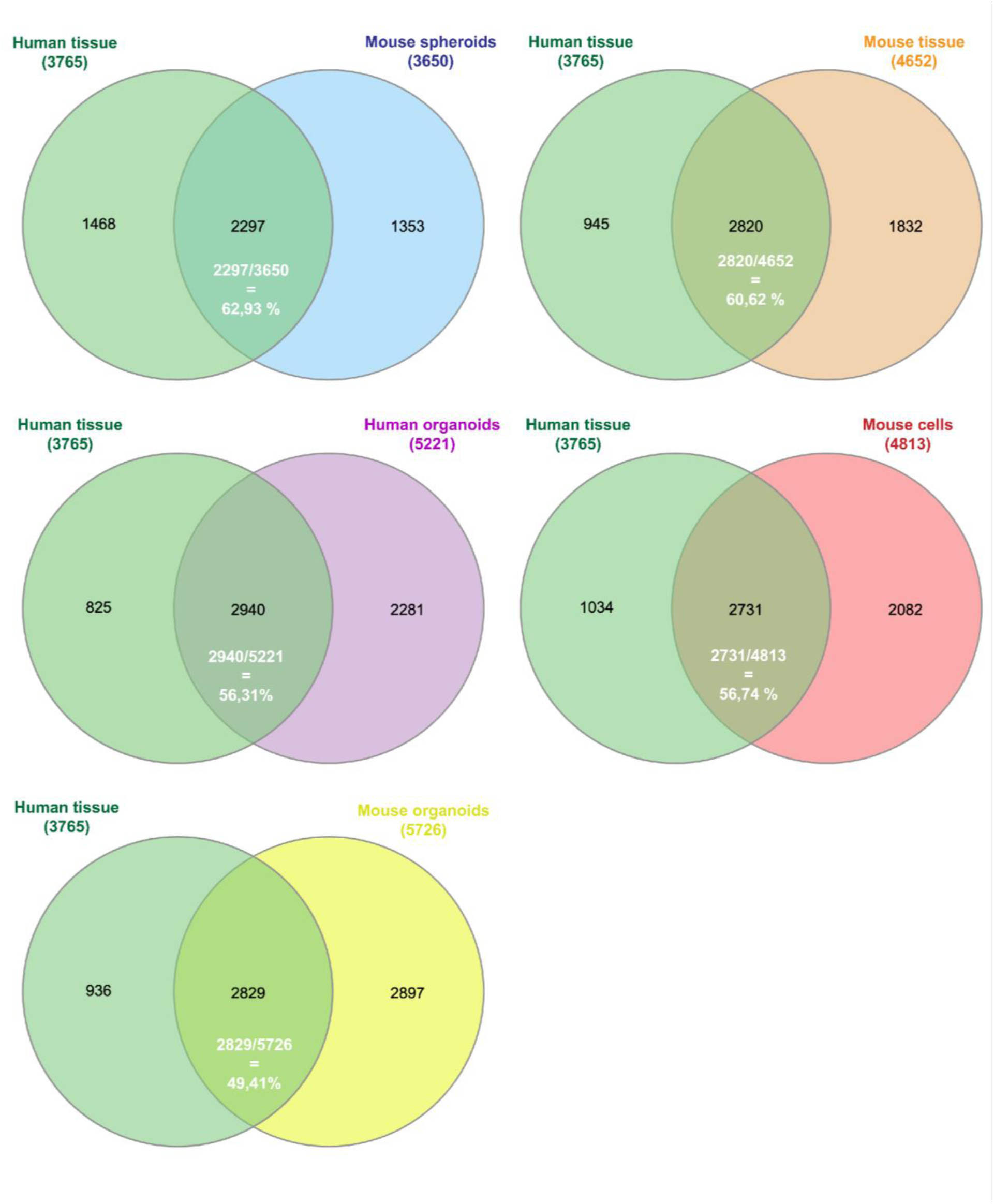
Venn diagrams comparing human PDAC with various PDAC research models. Note: Venn diagrams display the overlap of proteins identified in human PDAC tissue (n=3,765) with proteins detected in five different PDAC research models: mouse spheroids (n=3,650), mouse tissue (n=4,652), mouse cells (n=4,813), human organoids (n=5,221), and mouse organoids (n=5,726). The overlap percentages indicate the proportion of common proteins between human tissue and each research model, ranging from 49.41% (mouse organoids) to 62.93% (mouse spheroids).

A total of 1,975 proteins that were commonly expressed among all the six PDAC research models (Fig. 3). The proportion of these shared proteins varied among the different models, ranging from 35% to 54% (Table 1).

**Fig.3.**
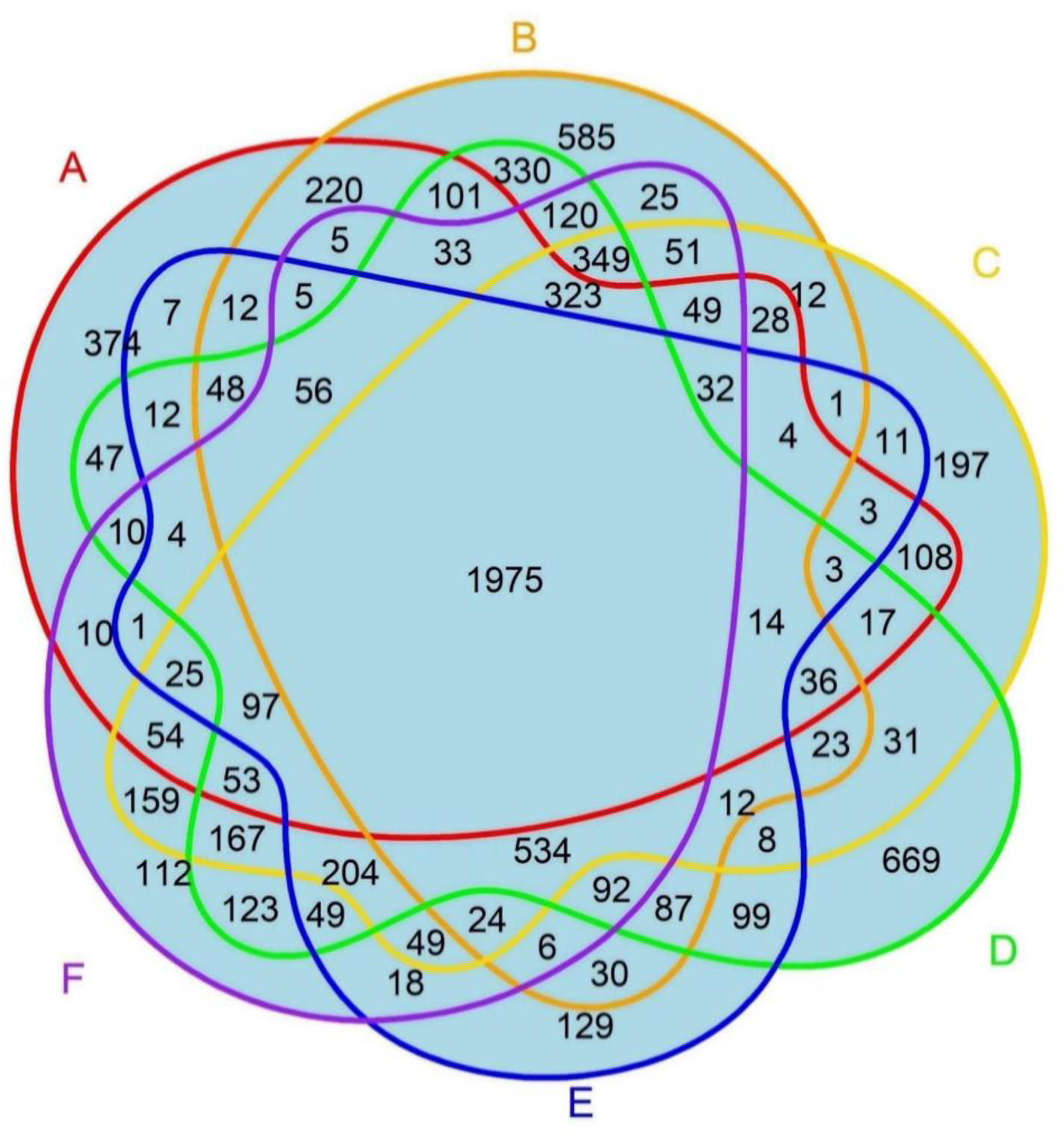
Venn diagram comparing six different PDAC models. Note: overlap of proteins identified among six PDAC groups: (A) Human PDAC tissue, (B) Human PDAC organoids, (C) Murine PDAC tissue, (D) Murine PDAC organoids, (E) Murine PDAC spheroids, and (F) Murine PDAC cells.

**Table 1:** Proportion of common proteins across research models.

| Research model | % of common proteins in research models |
| --- | --- |
| Murine PDAC cells | 1975/4813 = 41,0 % |
| Murine PDAC tumor tissue | 1975/4652 = 42,5 % |
| Murine PDAC spheroids | 1975/3650 = 54,1 % |
| Murine PDAC organoids | 1975/5726 = 34,5 % |
| Human PDAC organoids | 1975/5221 = 37,8 % |
| Human PDAC tumor tissue | 1975/3765 = 52,5 % |

### Knowledge presentation of hub proteins across various research models and human PDAC

PPI analysis of the 1,975 common PDAC proteins identified several hub proteins, including GAPDH, ACTB, HSP90AA1, HSPA8, HSP90AB1, HSPA4, EEF2, EFTUD2, RPS3, and HNRNPA1, which emerged as the top 10 hub proteins based on Cytoscape network analysis. Each of these proteins had 300 or more node connections. The 1,975 common proteins identified across all six PDAC research models were further analyzed for degree and betweenness centrality. Betweenness centrality was employed to assess a node’s significance by determining how frequently it acts as a bridge on the shortest path between two other nodes. ACTB showed the highest betweenness centrality and degree, followed by GAPDH, HSPA8, and HSP90AB1 (Fig. 4, and Table 2).

**Fig. 4.**
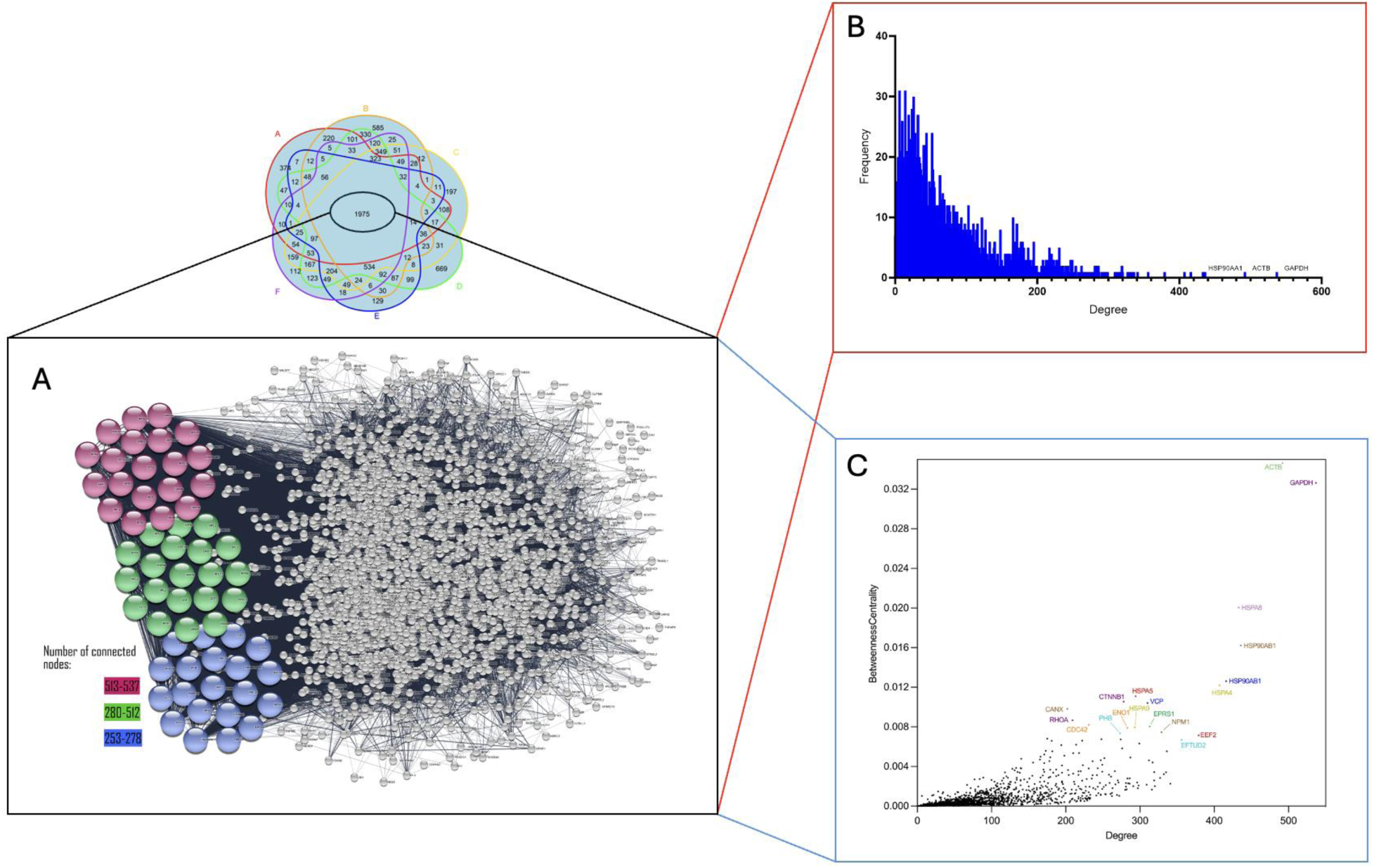
PPI of overlapped proteins (A), degree distribution (B) and betweenness centrality (C) across six research models. Note: the top 60 hub proteins are highlighted based on their degree of connectivity within the network. The top 20 proteins in purple, having 513-537 connective nodes, the next 21-40 proteins in green with 280-512 connections, and the proteins ranked 41-60 in blue with 253-278 connections (A). The degree distribution, where the x-axis represents the degree of connectivity (number of direct interactions) and the y-axis shows the frequency of each degree (B). Betweenness centrality is plotted against degree; the x-axis again represents the degree, while the y-axis measures betweenness centrality, indicating how often a protein acts as a bridge along the shortest path between two other proteins in the network (C). Each dot represents a protein, with ACTB exhibiting the highest betweenness centrality and degree, followed by GAPDH, HSPA8, and HSP90AB1.

**Table 2:** Top 10 hub proteins. The hub proteins are ranked by their degree of connectivity within the network. The proteins are highlighted with colors corresponding to the clusters depicted in Fig. 4A.

| Gene name | Degree | Betweenness Centrality | Description |
| --- | --- | --- | --- |
| HSP90A A1 | 436 | 0,01622 | Heat shock protein 90 alpha family class A member 1 |
| HSPA8 | 433 | 0,02005 | Heat shock protein family A (Hsp70) member 8 |
| HSP90A B1 | 416 | 0,01260 | Heat shock protein 90 alpha family class B member 1 |
| EEF2 | 379 | 0,00714 | Eukaryotic translation elongation factor 2 |
| VCP | 310 | 0,01041 | Valosin containing protein |
| RPL3 | 297 | 0,00179 | Ribosomal protein L3 |
| HSPA5 | 294 | 0,01109 | Heat shock protein family A (Hsp70) member 5 |
| HSPA9 | 293 | 0,00795 | Heat shock protein family A (Hsp70) member 9 |
| CTNNB1 | 278 | 0,01054 | Catenin beta 1 |
| PHB | 273 | 0,00736 | Prohibitin 1 |

### Knowledge presentation of hub canonical pathways

To gain deeper insights into PDAC progression and uncover potential early diagnostic biomarkers and therapeutic targets, hub canonical pathways were identified alongside hub proteins. Notably, several pathways with high betweenness centrality and a significant number of connections were highlighted, including the regulation of expression of SLITs and ROBOs, signaling of ROBO receptors, and NEP/NS2 interactions with cellular machinery. Additionally, key canonical pathways such as eukaryotic translation initiation, processing of capped intron-containing pre-mRNA, and SRP-dependent co-translational protein targeting to the membrane were identified (Fig. 5 and Tables 3 and 4).

**Fig. 5.**
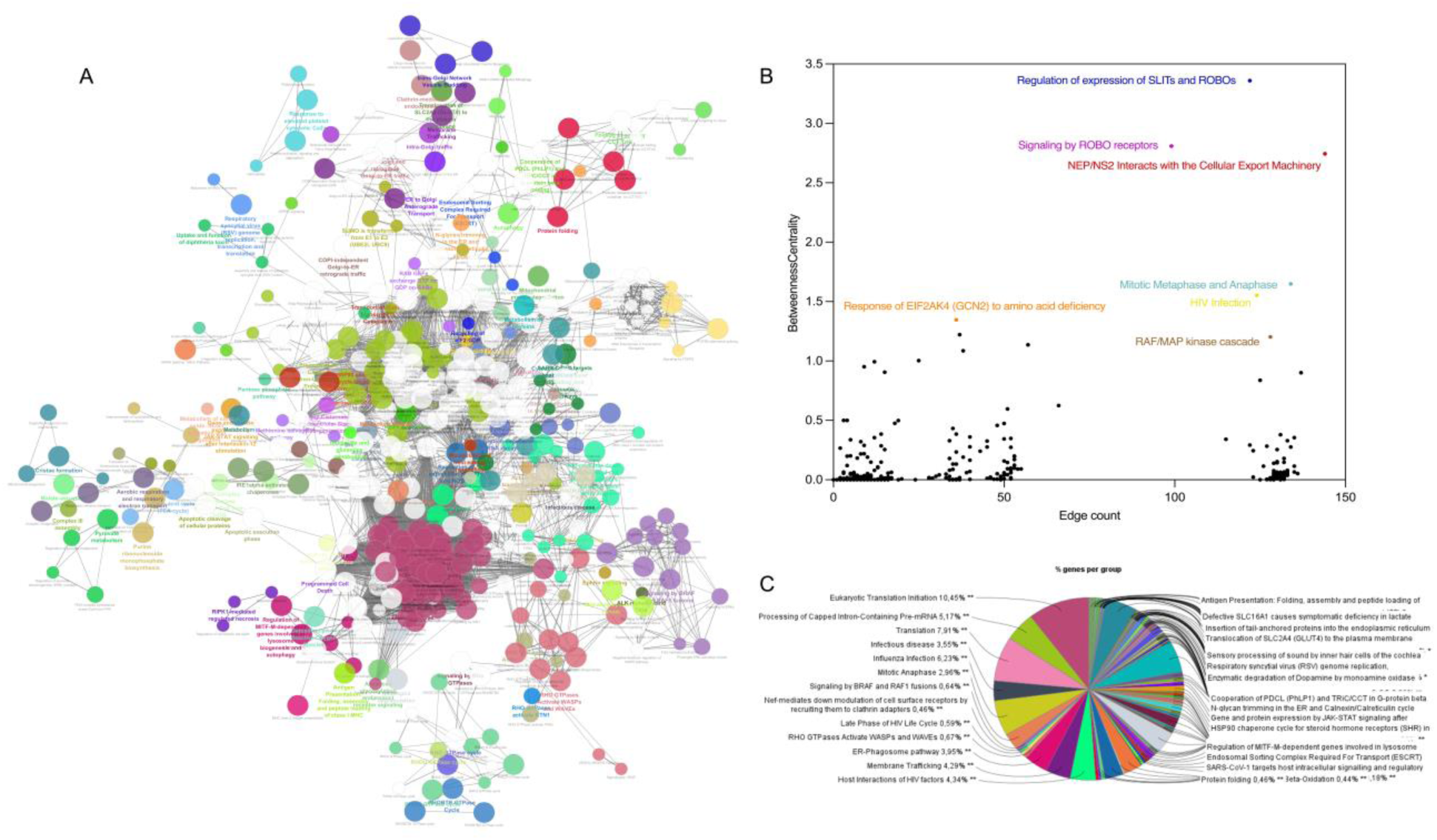
Network of hub canonical pathways (A), scatter plot of pathways in terms of betweenness centrality vs. edge count (B), and pie chart of distribution of genes per pathway (C). In A, different colors represent different pathways when a node (representing a gene or a pathway) is divided into multiple colors, because the genes associated with that node are involved in multiple pathways or functional groups. In B, each dot represents a pathway with x-axis in terms of edge count (degree centrality), and y-axis in terms of betweenness centrality. Pathways with higher edge counts are those that interact with more other pathways, while higher betweenness centrality values indicate pathways that serve as critical connectors or bridges within the network. The highlighted pathways represent those with the highest edge counts and significant betweenness centrality values, indicating their roles as critical hubs within the network. In C, each sector represents a specific pathway or biological process, with the size of the sector corresponding to the proportion of genes from the protein dataset involved in that pathway. **: p ≤ 0.05 after Benjamini-Hochberg correction.

**Table 3.** Connections of pathways sorted by betweenness centrality of edge counts. The edge count is the sum of indegree and outdegree. Here, the pathways are sorted on the betweenness centrality.

| Term | Betweenness Centrality | Edge Count | Indegree | Outdegree |
| --- | --- | --- | --- | --- |
| Regulation of expression of SLITs and ROBOs | 3,3599 | 122 | 15 | 107 |
| Signaling by ROBO receptors | 2,8086 | 99 | 65 | 34 |
| Disorders of transmembrane transporters | 2,7432 | 144 | 64 | 80 |
| Mitotic metaphase and anaphase | 1,6494 | 134 | 96 | 38 |
| Response of EIF2AK4 (GCN2) to amino acid deficiency | 1,3463 | 36 | 9 | 27 |
| Nuclear envelope (NE) reassembly | 1,2219 | 37 | 16 | 21 |
| RAF/MAP kinase cascade | 1,2031 | 128 | 58 | 70 |
| Nonsense-mediated decay (NMD) | 1,0862 | 38 | 10 | 28 |
| SARS-CoV-1-host interactions | 1,0032 | 25 | 8 | 17 |
| MAP2K and MAPK activation | 0,9931 | 12 | 8 | 4 |
| Post-translational protein modification | 0,9517 | 9 | 3 | 6 |
| Metabolism of RNA | 0,9059 | 15 | 2 | 13 |
| Deubiquitination | 0,8379 | 125 | 52 | 73 |
| Signal transduction by growth factor receptors | 0,6253 | 66 | 31 | 35 |
| Defective TPR towards thyroid papillary carcinoma | 0,5965 | 51 | 14 | 37 |
| Energy-dependent regulation of mTOR by LKB1-AMPK | 0,5000 | 4 | 1 | 3 |
| Protein localization | 0,5000 | 3 | 1 | 2 |
| SARS-CoV-2-host interactions | 0,4988 | 19 | 4 | 15 |
| SARS-CoV-1 Infection | 0,4758 | 18 | 8 | 10 |
| Mitotic prophase | 0,4741 | 50 | 11 | 39 |
| tRNA processing | 0,4574 | 48 | 6 | 42 |
| Metabolism of amino acids and derivatives | 0,4316 | 35 | 16 | 19 |
| Infectious disease | 0,3851 | 14 | 7 | 7 |
| Axon guidance | 0,3843 | 36 | 20 | 16 |
| Fc epsilon receptor (FCERI) signaling | 0,3788 | 131 | 97 | 34 |
| Transport of small molecules | 0,3603 | 40 | 30 | 10 |

**Table 4.**
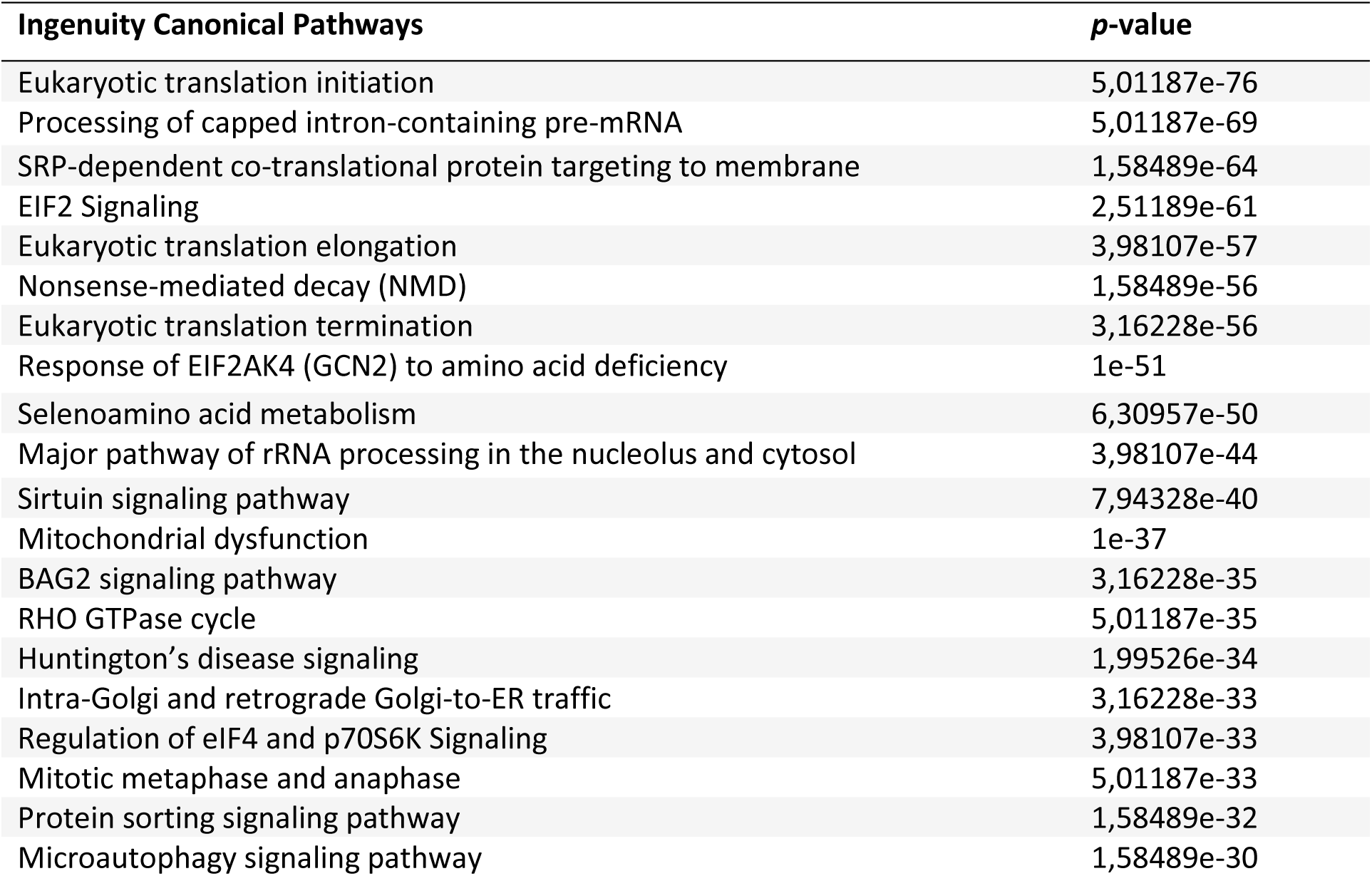
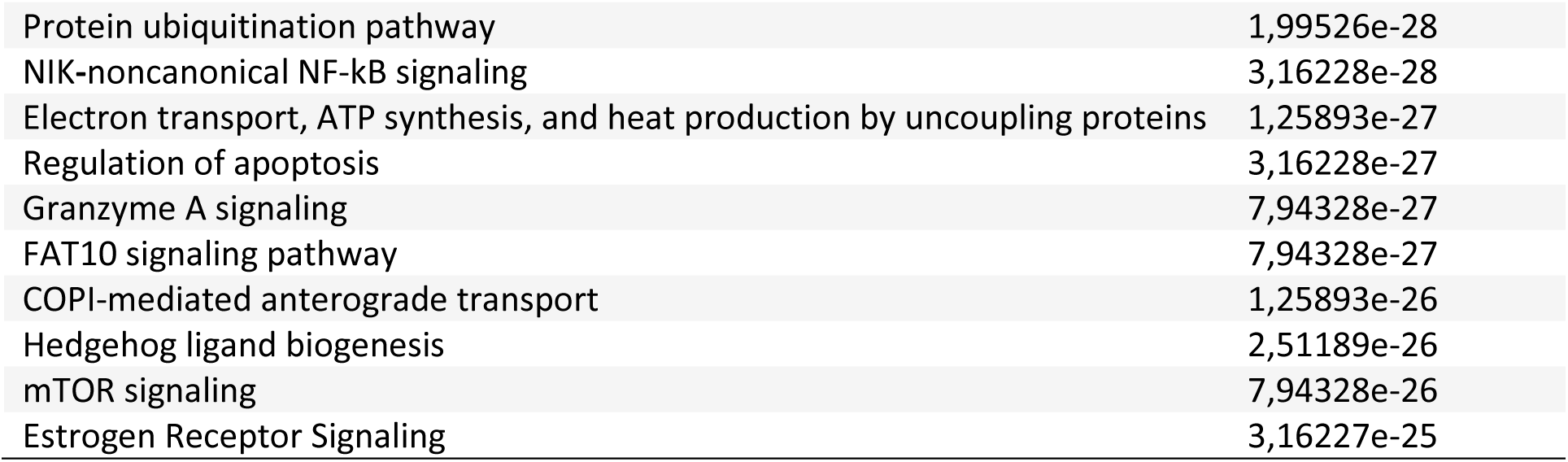
Top 30 canonical pathways based on 1,975 common proteins. The pathways are sorted based on *p*-value.

### Knowledge presentation of high topological proteins across various research models and human PDAC

The topological coefficient is a relative measure for the extent to which a node shares neighbors with other nodes. Nine AI/ML models were used and compared to determine which model best captured the relationship between node degree and topological coefficient in the network dataset consisting of the 1,975 common PDAC proteins (Fig. 6). Each model was trained using 5-fold cross-validation, and its predictive performance was assessed using three key evaluation metrics. R² Score (Coefficient of Determination) shows how well the model explains variance in the data. Mean Absolute Error (MAE) represents the average magnitude of prediction errors. Mean Squared Error (MSE) is similar to MAE but penalizes larger errors more heavily. Among the models tested, tree-based ensemble methods consistently outperformed other regression approaches. The Random Forest Regressor achieved the highest R² score (0.7668) and the lowest MAE (0.0220), indicating that it was the most reliable model for predicting the topological coefficient. The Gradient Boosting Regressor (R² = 0.7589, MAE = 0.0225) and CatBoost Regressor (R² = 0.7591, MAE = 0.0224) performed nearly as well. This observation suggests that boosting-based AI/ML approaches are highly effective in modeling trends for topological coefficient. The performance differences between these models were marginal. This implies that any of these three models could serve as viable options for regression analyses on similar network datasets. The Gaussian Process Regressor (R² = 0.7152, MAE = 0.0246) also provided a reasonable fit to the data but exhibited some instability, particularly at higher degree values, due to observed oscillations in predictions. This model captured nonlinear variations well but was more sensitive to noise in the dataset.

**Fig. 6.**
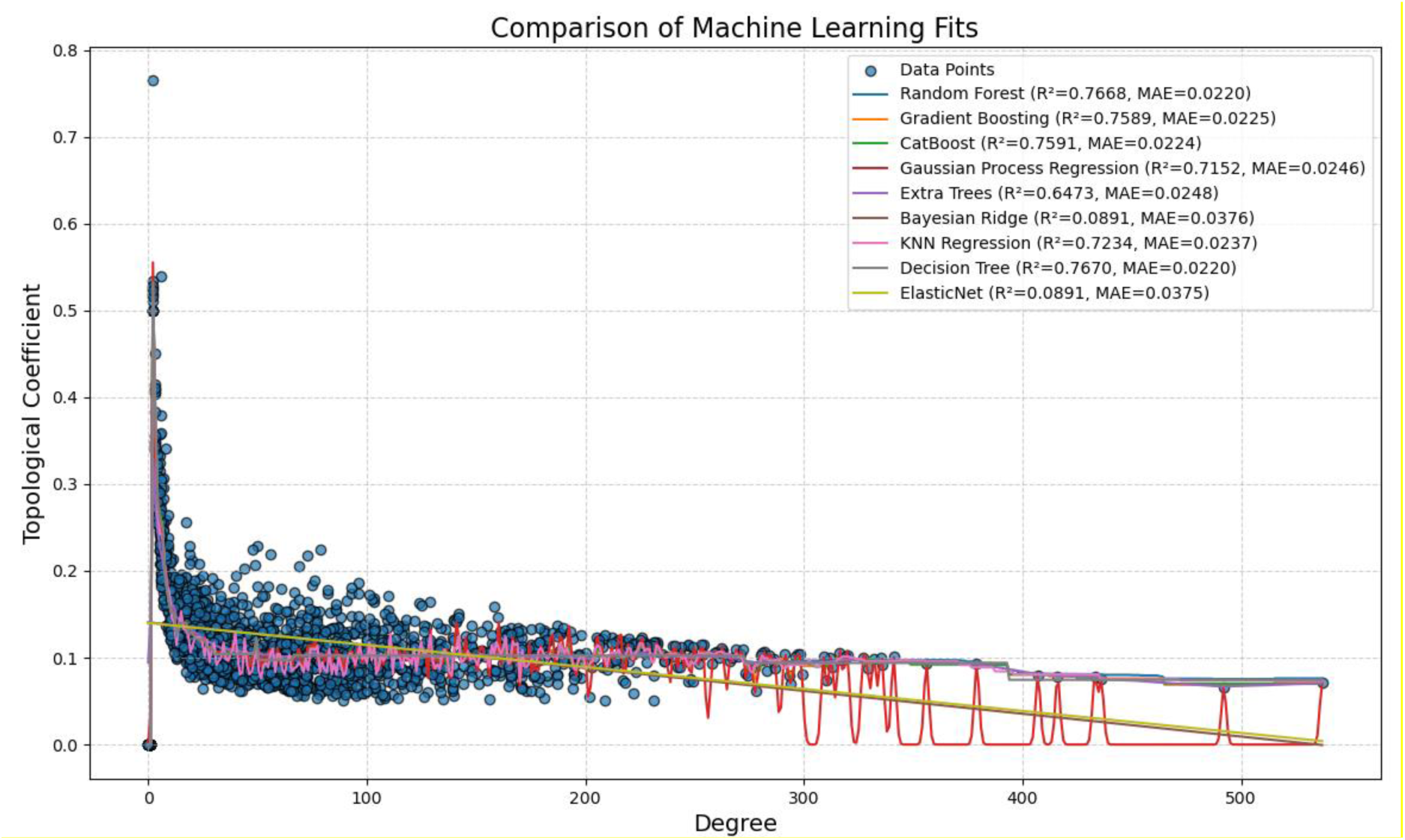
Scatter plot of topological coefficient against degree. Note: overlaid curves of various regression models including tree-based ensemble methods (Random Forest, Gradient Boosting, CatBoost, Extra Trees, and Decision Tree), probabilistic approaches (Gaussian Process Regression), distance-based methods (K-Nearest Neighbors), and linear regularization models (Bayesian Ridge and ElasticNet). Performance metrics, including R² (explained variance) and Mean Absolute Error (MAE), for each model are displayed in different colors.

Machine leaning with Random Forest, Gradient Boosting, and CatBoost showed the best predictive accuracy, capturing the nonlinear relationship between degree and topological coefficient with minimal error. Comparably, linear models such as Bayesian Ridge and ElasticNet failed to capture the underlying trend, while Gaussian Process Regression exhibited instability at higher degrees (Fig. 6). Of note, the distribution curves showed most nodes with coefficient more than zero and an inverse relationship the coefficient and degree (Fig. 6 and Table 5).

**Table 5:**
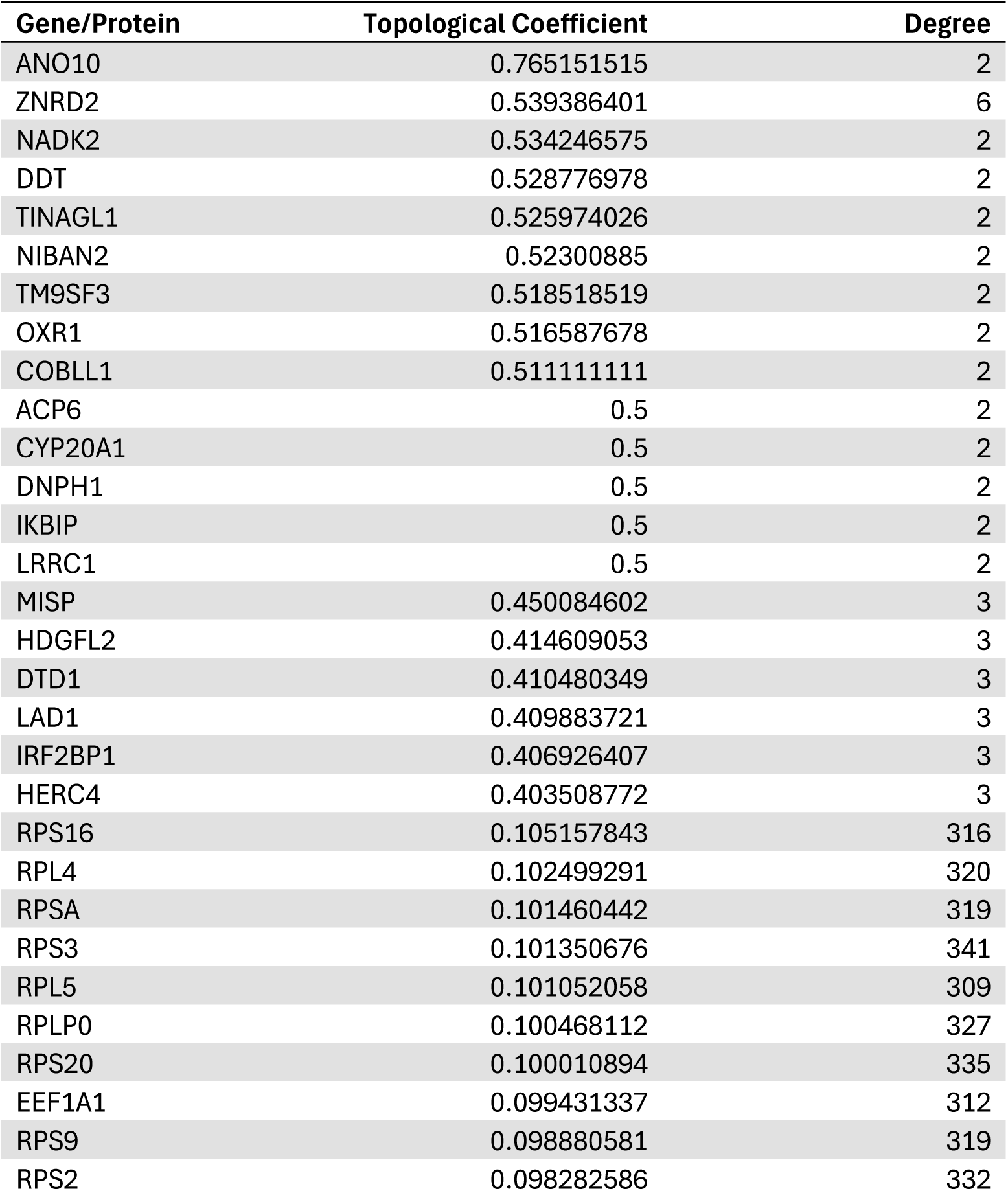
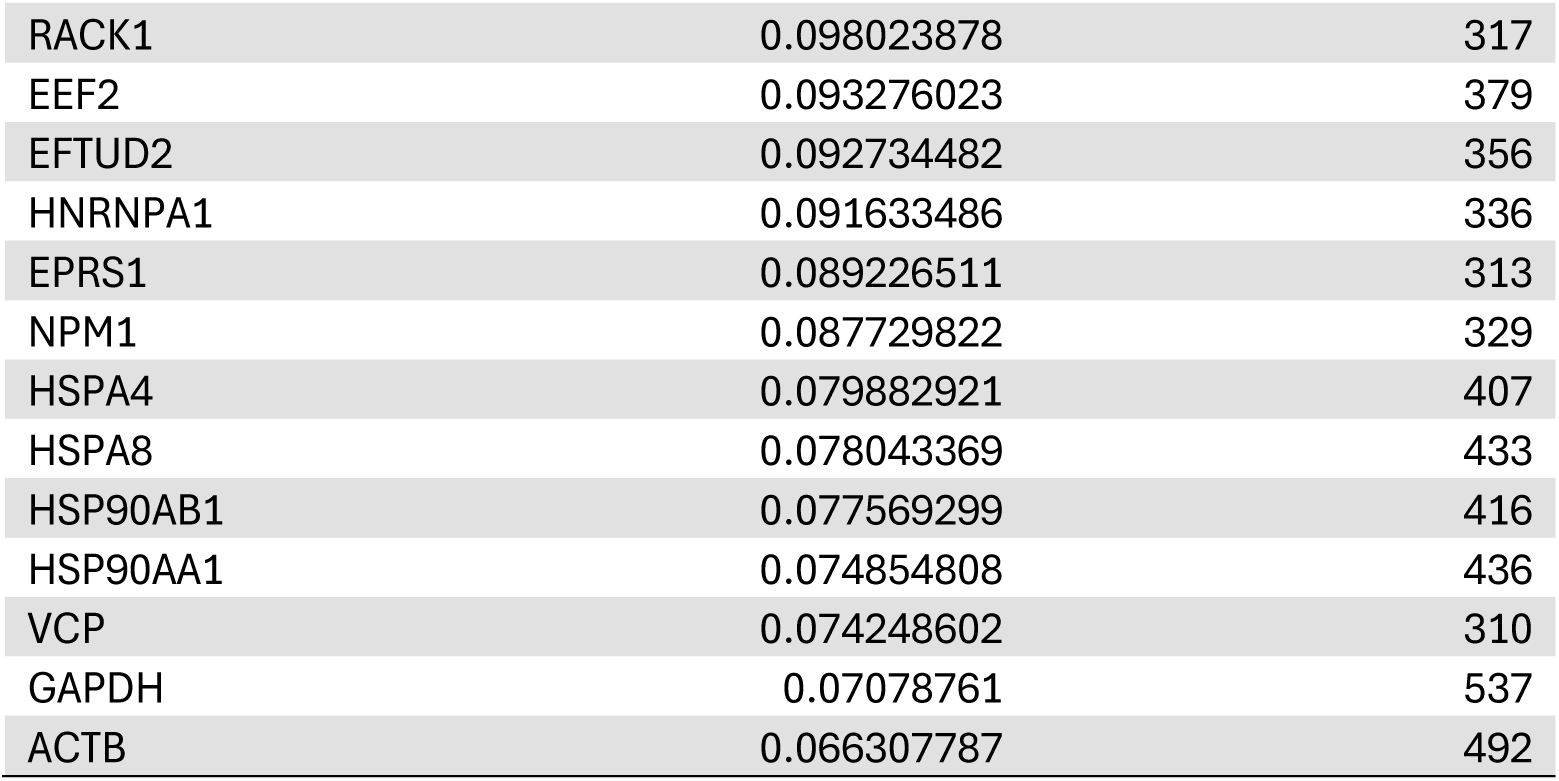
Topological coefficients and degree values for nodes representing a single gene/protein. Data cutoff at topological coefficient ≥ 0.4 and degree ≥ 300. High topological coefficient coincides with low degree and high degree coincides with low topological coefficient.

| Gene/Protein | Topological Coefficient | Degree |
| --- | --- | --- |
| ANO10 | 0.765151515 | 2 |
| ZNRD2 | 0.539386401 | 6 |
| NADK2 | 0.534246575 | 2 |
| DDT | 0.528776978 | 2 |
| TINAGL1 | 0.525974026 | 2 |
| NIBAN2 | 0.52300885 | 2 |
| TM9SF3 | 0.518518519 | 2 |
| OXR1 | 0.516587678 | 2 |
| COBLL1 | 0.511111111 | 2 |
| ACP6 | 0.5 | 2 |
| CYP20A1 | 0.5 | 2 |
| DNPH1 | 0.5 | 2 |
| IKBIP | 0.5 | 2 |
| LRRC1 | 0.5 | 2 |
| MISP | 0.450084602 | 3 |
| HDGFL2 | 0.414609053 | 3 |
| DTD1 | 0.410480349 | 3 |
| LAD1 | 0.409883721 | 3 |
| IRF2BP1 | 0.406926407 | 3 |
| HERC4 | 0.403508772 | 3 |
| RPS16 | 0.105157843 | 316 |
| RPL4 | 0.102499291 | 320 |
| RPSA | 0.101460442 | 319 |
| RPS3 | 0.101350676 | 341 |
| RPL5 | 0.101052058 | 309 |
| RPLP0 | 0.100468112 | 327 |
| RPS20 | 0.100010894 | 335 |
| EEF1A1 | 0.099431337 | 312 |
| RPS9 | 0.098880581 | 319 |
| RPS2 | 0.098282586 | 332 |
| RACK1 | 0.098023878 | 317 |
| EEF2 | 0.093276023 | 379 |
| EFTUD2 | 0.092734482 | 356 |
| HNRNPA1 | 0.091633486 | 336 |
| EPRS1 | 0.089226511 | 313 |
| NPM1 | 0.087729822 | 329 |
| HSPA4 | 0.079882921 | 407 |
| HSPA8 | 0.078043369 | 433 |
| HSP90AB1 | 0.077569299 | 416 |
| HSP90AA1 | 0.074854808 | 436 |
| VCP | 0.074248602 | 310 |
| GAPDH | 0.07078761 | 537 |
| ACTB | 0.066307787 | 492 |

Several other models were tested, showing a moderate predictive ability. The Extra Trees Regressor (R² = 0.6473, MAE = 0.0248) introduces additional randomness to tree splitting, but performed worse than Random Forest. This suggests that the increased variance introduced by Extra Trees did not lead to improved generalization in this case. The K-Nearest Neighbors (KNN) Regressor (R² = 0.7234, MAE = 0.0237) showed surprising effectiveness, likely due to its ability to capture local patterns in the dataset. However, it struggled to generalize globally across the entire degree range, which limits its applicability for large-scale network predictions.

Among the models that underperformed, the Decision Tree Regressor (R² = 0.7670, MAE = 0.0220) was competitive in terms of R² and MAE, but its predictions exhibited overfitting. This resulted in abrupt changes in predicted values across different degree values. In contrast, Bayesian Ridge Regression (R² = 0.0891, MAE = 0.0376) and ElasticNet Regression (R² = 0.0891, MAE = 0.0375) failed to capture meaningful relationships, as they assume a linear dependency between degree and topological coefficient, which does not align with the true nonlinear nature of the dataset.

### Knowledge presentation of PDAC signaling network ‘signature’

It is a common idea to identify specific biomarkers that are expressed only by PDAC cancerous cells. By the systems modeling further filed with normal controls (i.e., normal mouse pancreas and normal mouse exocrine organoids), only 15 proteins were shown up (Fig. 7A). Based on TCGA dataset at Human Protein Atlas, the following were of low cancer specificity, including FAHD1, NEK9, SURF1, RHOC, LAMTOR2, TMED3, ACOT13, RAB1A, EPHB4, SCO1, MYO18, and BABAM1, whereas TOMM6 was not detected in cancer and SERPINB9 was highly expressed in testis cancer, respectively. Furthermore, 74% (i.e., 1,462 out of 1,975) PDAC proteins were also present in normal tissue, probably indicating that the ‘building blocks’ were similar but the ‘building appearance and function as “signature in terms of the signaling networks” differ between PDAC and the normal tissue (Fig. 7B and 7C). Significant pathways of the PDAC proteins (‘1462’) in pie chart included Cap-dependent translation initiation, eukaryotic translation initiation, and processing of capped intron-containing pre-mRNA (Fig. 7B), whereas significant pathways of the non-neoplastic proteins (‘813’) in pie chart included platelet sensitization by LDL, integrin signaling, cell cycle, metabolism of RNA, laminin interactions, and MASTL facilitates mitotic progression (Fig. 7C). It should be noticed in terms of betweenness centrality *vs* edges that 73 of these pathways were shared by the PDAC and the non-neoplastic proteins, such as metabolism of RNA, mitotic prophase, M phase, signal transduction by growth factor receptor, mitotic metaphase and anaphase and nuclear envelope reassembly (Fig. 7D and 7E).

**Fig. 7.**
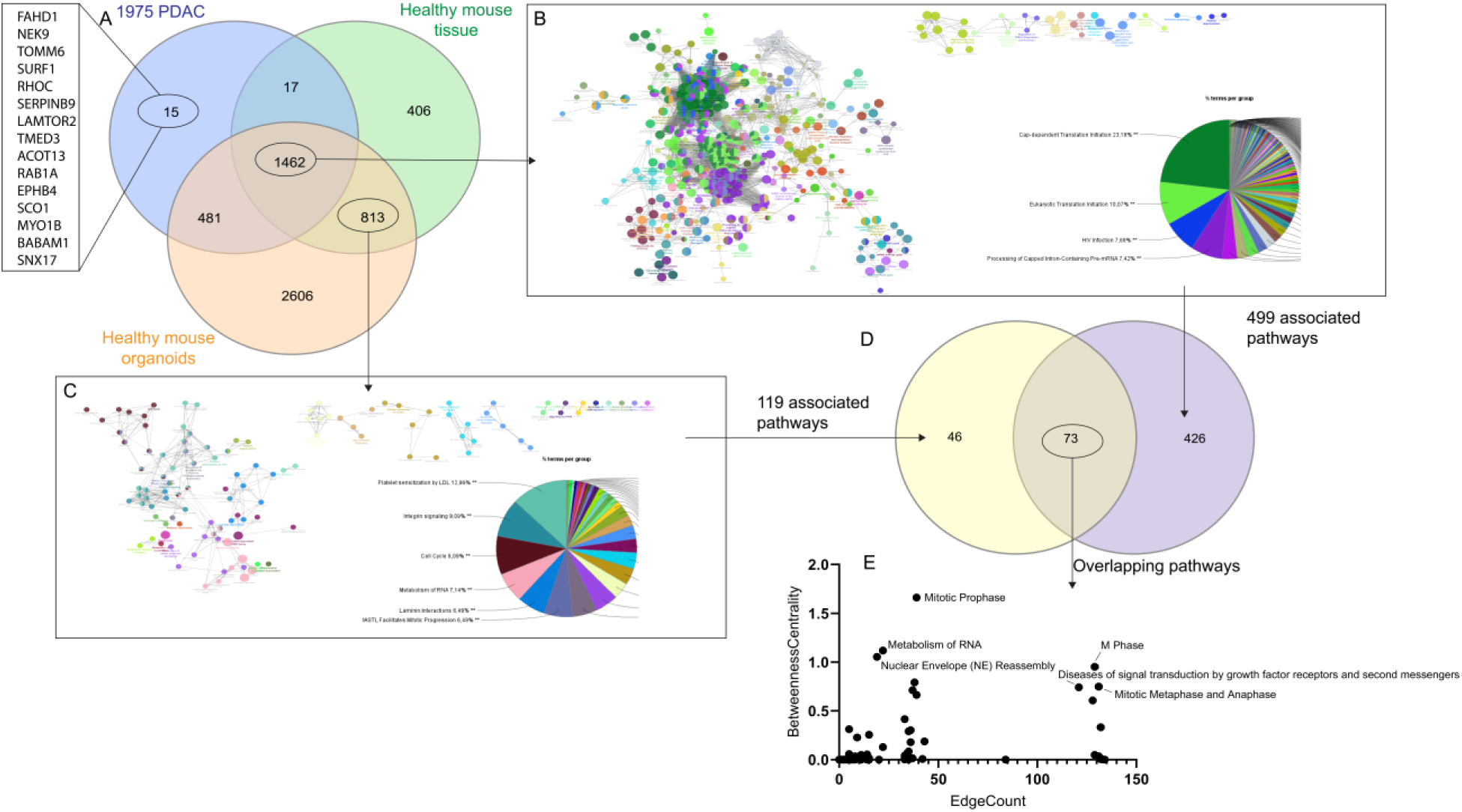
Comparative analysis of PDAC and non-cancerous models with pathway associations. Venn diagram depicting the 1,975 common PDAC proteins and their overlap with proteins from healthy mouse pancreatic tissue and healthy mouse pancreatic organoids. The unique proteins for PDAC are listed in the left column (A). Visualization of the pathways associated with the proteins common to all groups in the Venn diagram (i.e., overlapping proteins between PDAC models and healthy models). The pathways are color-coded, and the pie chart provides a breakdown of the pathway categories (B). Pathways associated exclusively with proteins found in healthy models (non-cancerous) (C). Venn diagram showing the overlap between pathways associated with healthy models and cancerous PDAC models. This highlights the shared pathways. Scatter plot illustrates the relationship between edge count and betweenness centrality for the identified pathways (E).

## Discussion

In the field of translational research, ‘systems biology’ is the state-of-the-art approach using multi-omics, such as transcriptomics and proteomics to acquire data for constructing models of complex biological systems and diseases^13,14^. However, extracting high-quality RNA from pancreatic tissue, including pancreatic tumors, poses significant technical challenges for many laboratories due to the high ribonuclease content in the pancreas ^15–17^. It should also be noticed that discrepancies between mRNA expression and protein abundance are common, especially in the pancreas^18,19^. Thus, the datasets used in the present study were proteomics that were standardized and annotated from various PDAC research models and patient samples^1^. The results of the systems modeling showed the knowledge presentations, which led to the generation of hypothesis that the common proteins among the PDAC research models and patient samples, particularly hub proteins, hub canonical signaling pathways, high topological proteins and signature pathways, are the targets to overcome the ‘translational gap’ (Fig. 8).

**Fig. 8.**
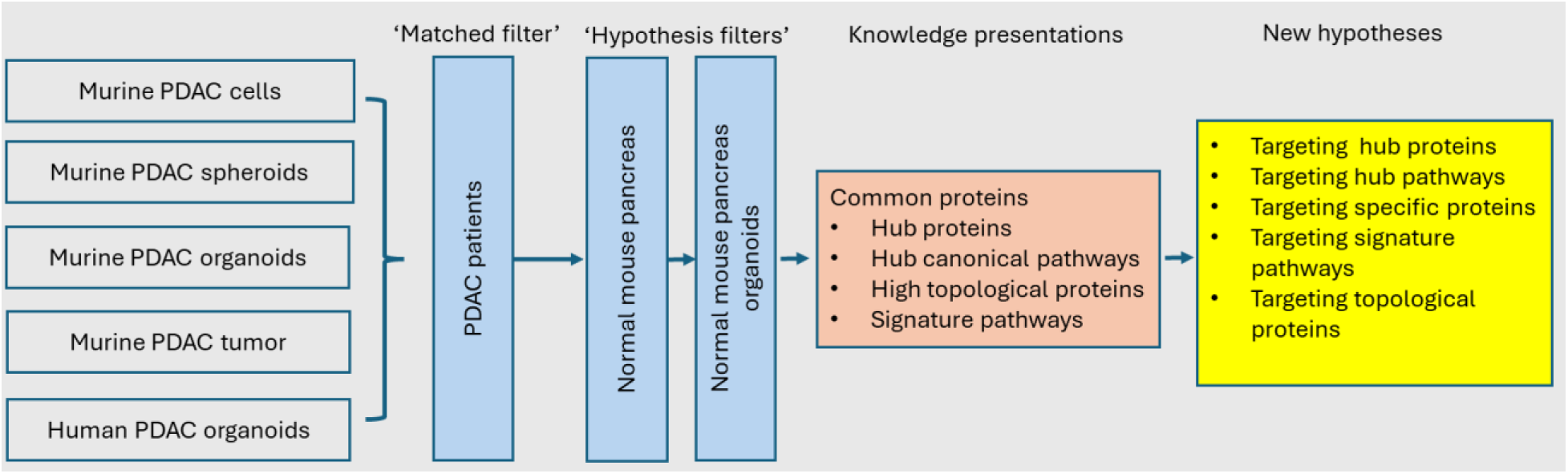
Summary of main results. Nore: study models, filters, knowledge presentations and new hypotheses for the translational research of PDAC.

### Targeting common proteins, particularly hub proteins

The results of the present study showed 50-60% of proteins being common across these models between PDAC models and human PDAC tissues. The hub proteins, such as CCT4, EZR, and GAPDH, are known to be involved in critical pathways like neutrophil degranulation and sirtuin signaling, which are linked to the altered metabolic profiles and increased cancer stemness observed in neoadjuvant-treated PDAC. PPI analysis revealed that the distribution of protein interactions followed an inverse power law, indicative of a scale-free network topology, where correlations between degree and betweenness (“bottleneck-ness”) centrality confirmed the flow of communication within the network. Nodes with both high degree and high betweenness centrality are particularly significant as they lie on key communication paths and can regulate the flow of information within the network ^20^. A comparative study in yeast identified a core set of 2,708 proteins with an average of 5.26 PPIs per protein. In the present study involving 1,975 proteins, the average number of PPIs per protein was 75.6 with the top 60 proteins exhibiting 253-537 PPIs each, reflecting the complexity of the PDAC network. As expected, housekeeping gene-encoded proteins, such as GAPDH and β-actin, emerged as top hub proteins that are crucial for cellular metabolism, homeostasis, motility, structure, and integrity. However, it would be impossible to target ACTB unless its specific isoforms for PDAC are identified. The next set of top hub proteins included heat shock proteins (HSPs), vital for maintaining cellular proteostasis through the integration of protein folding and degradation processes^21^. Notably, β-catenin, a well-known protein under investigation as a potential target in cancer treatment, particularly in PDAC, also ranked among the top hub proteins ^22,23^. Ribosomal proteins, essential for the translation process and indicative of the high metabolic and proliferative demands of PDAC cells, also featured prominently among the hub proteins. When considering potential therapeutic targets, it is important to recognize that both the hub protein itself and its interaction partners must be specific in order to avoid off-target effects.

### Targeting hub pathways

The present study showed the hub canonical pathways defined by hub proteins with high betweenness centrality and high edge count. For instance, regulation of expression of SLITs and ROBOs and signaling of ROBO receptors might represent new targets^24^. NEP/NS2 interacts with cellular machinery is known to be involved with the influenza virus NEP (NS2 protein)^25^, and has not been reported in connection with cancer, particularly PDAC (until this study). The present study also identified the hub canonical pathways defined by the percentages of its hub proteins included eukaryotic translation Initiation ^26^, processing of capped intron-containing pre-mRNA ^27^, and SRP-dependent co-translational protein targeting to membrane ^28,29^.

Several additional highly connected pathways were identified, including the RAF/MAP kinase cascade, mitotic metaphase and anaphase, and response of EIF2AK4 (GCN2) to amino acid deficiency, all of which are relevant to cancer progression and therapy resistance. The RAF/MAP kinase cascade is a well-established driver of PDAC proliferation and survival, frequently altered in pancreatic tumors. Similarly, mitotic metaphase and anaphase regulation is crucial for tumor cell division and chromosomal stability, making it a potential therapeutic target. The EIF2AK4 pathway, involved in the cellular response to amino acid deprivation and metabolic stress, has been linked to tumor adaptation under nutrient-limited conditions, which is particularly relevant for PDAC’s hypoxic and nutrient-deprived microenvironment. The identification of these pathways supports their potential roles in tumor progression and resistance mechanisms in PDAC and highlights key regulatory networks that may be further explored for therapeutic targeting.

### Targeting specific proteins

Despite of hard attempts during the past several decades, no specific proteins or protein signatures have been conclusively identified as reliable clinical indicators for risk assessment, early detection, progression, treatment response, or prognosis for PDAC. While the 15 proteins identified in the present study are unlikely to serve as definitive PDAC-specific biomarkers, the broader set of 1,462 proteins may hold potential as PDAC-unique proteins, particularly when considering personalized medicine approaches, such as filtering with individual patient proteomic data. Additionally, the 813 ‘normal’ proteins, identified by filtering against normal mouse pancreas and pancreatic organoids, could be further refined using normal human pancreatic protein data. The systems modeling approach is adaptable, allowing for various types of ‘filtering’ - for example, isolating cancerous cells from the tumor to exclude contributions from the tumor microenvironment, including immune cells. It is believed that non-neoplastic cells within the tumor, such as those associated with chronic pancreatitis, may increase the risk of PDAC. The findings of the present study may inspire further research into identifying PDAC initiators or promoters beyond inflammation, such as cytokine interleukin 33 ^30^.

### Targeting specific pathways

The ‘PDAC signaling network signature’ proposed in the present study highlights the pivotal roles of signaling pathways associated with translation initiation, including cap-dependent translation initiation, eukaryotic translation initiation, and the processing of capped intron-containing pre-mRNA^31–35^. These pathways are central to the functionality of PDAC cells, reflecting their high translational activity and protein synthesis demands. Additionally, the significant pathways identified in non-neoplastic proteins may be viewed as key contributors to the ‘PDAC tumor microenvironment’ excluding immune cells. These pathways include metabolism of RNA, platelet sensitization by LDL, integrin signaling, cell cycle regulation, laminin interactions, and MASTL-facilitated mitotic progression^36–42^. Moreover, potential communication pathways between cancerous cells and tumor microenvironment cells appear to be active during mitotic phases, including prophase, M phase, metaphase, and anaphase, as well as during nuclear envelope reassembly. Diseases of signal transduction involving growth factor receptors and second messengers further underscore the complexity of the interactions within the PDAC microenvironment ^43^.

### Targeting topological proteins

Proteins with high topological coefficients in a network are sometimes defined as key nodes that are similar to the hubs in a network structure. The key nodes can directly determine the robustness and stability of the network. By effectively identifying and protecting these critical nodes, the robustness of the network can be improved, making it more resistant to external interference and attacks^44^. There is little literature on the top key nodes in connection with PDAC yet, which is worthful for further investigation, e.g. transmembrane 9 superfamily member 3 (TM9SF3) protein ^45^ and mitotic spindle positioning (MISP) protein which is an actin-bunding protein.

### Notes on AI/ML

The results of the present study showed that the tree-based models (Random Forest, Gradient Boosting, and CatBoost) produced smooth well-aligned curves that closely followed the distribution of actual data points. Their ability to learn from the underlying structure of the dataset resulted in highly accurate predictions with minimal deviation from observed values ^46–48^. In comparison, the Gaussian Process Regressor displayed high variability, specifically in regions where data density was low ^49^. The Decision Tree Regressor generated highly fragmented predictions, further highlighting its tendency to overfit individual training points rather than generalizing globally across the dataset ^50^.

Linear models, including Bayesian Ridge and ElasticNet, produced nearly flat trend lines with little predictive power ^51,52^. This implies that a simple linear dependency is inadequate for modeling topological coefficients in biological networks. Given the forementioned findings, Random Forest emerges as the most effective model, with Gradient Boosting and CatBoost as strong alternatives. These results suggested that ensemble tree-based methods are optimal for modeling the topological coefficient in network analysis, as they effectively handle nonlinearity and complex interactions between degree and connectivity patterns.

In conclusion, the systems modeling workflow developed in the present study may provide a robust framework for bridging the ‘translational gap’ by identifying potential ‘matched targets.’ The findings, along with the predictions and hypotheses generated, warrant further validation through additional models. Moreover, this approach supports the refinement of existing hypotheses and the data presented in connection with the present study may open avenues for generating new hypotheses related to PDAC initiation, early detection, and prevention strategies.

## Methods

### Ethics

Raw data from human participants and animals were collected in accordance with the ethical standards of the institutional and/or national research committee and the Helsinki Declaration and its later amendments or comparable ethical standards (for humans: REK South-east 2015/738; for animals: Norwegian Food Safety Authority Mattilsynet FOTS 8335).

### Data processing

The information about the datasets used in this study can be found in our recently published paper entitled “Proteomics profiling of research models for studying pancreatic ductal adenocarcinoma” (DOI: 10.1038/s41597-025-04522-x). This publication details the preparation, validation, and proteomic profiling of the research models, ensuring transparency and reproducibility. Aspects such as sample collection, proteomics analysis using LC-MS/MS, and data processing, including quality control, were performed following the protocols outlined in that work. The corresponding proteomic datasets generated in the study are publicly available through the PRIDE database under the accession numbers PXD057793, PXD057795, PXD057798, PXD057804, PXD057888, PXD057829, PXD057928, and PXD057607.

To ensure high-confidence protein identification, MS data were processed using MaxQuant (version 2.3.1.0) with standard filtering steps, including removal of contaminants, reverse proteins, and proteins identified only by site modifications. Within each model, proteins were considered reliably detected if they appeared in at least 50% of biological replicates (e.g., identified in at least 3 out of 6 replicates for models with six replicates). This filtering step reduced variability and ensured robust protein detection within each model.

### Bioinformatic analysis

RStudio (ggVennDiagram-package in Rstudio), Cytoscape 3.10, Ingenuity Pathways Analysis (IPA). ClueGO analysis of the 1975 common proteins was performed in Cytoscape v3.9.1 (ClueGO v2.5.10) using Homo sapiens (NCBI Taxonomy ID: 9606) and Reactome pathways (version 20.06.2024) as the reference set. Gene enrichment was assessed using a right-sided hypergeometric test with Benjamini-Hochberg correction and a p-value cutoff of 0.05, while GO levels were restricted to 5–11 to balance specificity. Pathways were clustered using a Kappa score threshold of 0.4, which optimizes functional grouping by balancing sensitivity and specificity. The initial group size of 1 and 50% sharing ratio were chosen based on ClueGO’s iterative algorithm, which first identified 180 groups and then merged redundant clusters in two iterations, yielding 121 final pathway groups. This ensured biologically meaningful pathway associations while minimizing weak overlaps. After applying significance selection criteria, 566 representative terms remained, covering 1597 genes (80.86%), while 340 genes (17.22%) lacked functional annotations. The final network visualization in Cytoscape provided insights into the key functional pathways enriched in the dataset

The Kappa score in ClueGO is used to determine the functional similarity between pathways based on shared genes. It measures how much gene overlap exists between different pathways while adjusting for the likelihood of random matches. The default threshold of 0.4 is chosen to balance specificity and sensitivity, ensuring that only functionally related pathways are grouped together while minimizing noise from weak associations. This threshold is commonly used in bioinformatics tools employing Kappa statistics, where a cutoff of ≥0.35 was found to effectively distinguish meaningful biological relationships from random associations^53^. In ClueGO, pathways exceeding the chosen threshold are clustered together into functionally coherent groups, facilitating biological interpretation in a network-based visualization.

IPA uses the Right-tailed Fisher’s Exact Test to determine which pathways that are significantly enriched in the dataset. The molecules in the uploaded dataset were compared to a reference set. The null hypothesis (H0) is explained as the overlap of molecules for a particular biological category is due to chance alone. Any results *p*<0.05 allowed rejection of the null hypothesis. The Fisher exact test was used to ask whether the null hypothesis could be rejected and instead said that the overlap was larger than what would be expected by random sampling of the dataset. The significance level was only placed in the right tail of the distribution rather than in both tails. This was because there was interest to find whether there were more rather than fewer overlapping molecules associated with a particular pathway.

### AI/ML

AI was applied as a method to generate python scripts, using ChatGPT (GPT 4o) (Large Language Model, retrieved from https://openai.com/chatgpt/). It facilitated rapid debugging and iterative refinement, allowing for efficient implementation of complex ML pipelines (see below). To ensure robust evaluation, each model was trained and validated to allow for better generalization by reducing bias and variance. The AI-assisted scripting was used to automatically split the dataset, train models, and compute evaluation metrics, significantly reducing the need for time consuming manual labor. These metrics were calculated automatically for each model, with AI-assisted programming generating performance summaries. This ensured the identification of the best predictive model and python was used to generate scatter plots with ML fitted curves for each model. The visualization provided a comparative analysis of model performance, making it possible to assess how well each model captured the relationship between degree and topological coefficient.

#### ML preprocessing

The dataset used in the regression analysis was obtained from a Cytoscape network analysis ^54^ made on the common 1975 proteins, where each node represents a protein, and edges indicate interactions between them. Each node in the network can be characterized by degree, which represents the number of connections it has, as well as topological coefficient, which is a measure of the similarity between a node’s connectivity pattern and that of its neighbors. The aim of the analysis was to investigate the topological coefficient based on the degree of each node. The reason being degree as a fundamental centrality measure in network science and often correlates with a node’s functional importance. Given the complex and nonlinear nature of biological networks, traditional linear models may not properly capture the relationships between these variables, necessitating the use of AI/ML approaches. Data preprocessing involved extracting relevant information from the network-based dataset, specifically the node degree as the predictor variable (x) and topological coefficient as the target variable (y). No additional scaling of the input data was performed, as many tree-based models inherently handle input data scaling and normalization. The dataset was used in its entirety without filtering or transformation to allow models to learn from the full distribution of degree values. 5-fold cross-validation was applied to evaluate each model’s predictive performance. This method splits the dataset into five equal parts, using four for training and one for testing. Subsequently, it iterates the process across all five subsets to obtain an average performance score.

#### Evaluation

A diverse set of AI/ML regression models was applied to predict the topological coefficient from node degree. The models included tree-based ensemble methods, which are highly effective for capturing nonlinearity in structured datasets. They also included non-tree-based methods, which serve as baseline comparisons for evaluating alternative approaches. Among the tree-based ensemble methods, Random Forest ^46–48^ and Extra Trees ^55^ were implemented. These models are known for their ability to capture nonlinear relationships through recursive partitioning. They use multiple decision trees to improve prediction accuracy, with Random Forest and Extra Trees averaging multiple trees to reduce variance, while Gradient Boosting and CatBoost refine predictions to minimize errors. Gaussian Process Regression (GPR)^49^ was included as a probabilistic approach that models the data using a kernel function to capture uncertainty and complex relationships between degree and topological coefficient. While highly flexible, GPR can sometimes be sensitive to noise and computationally expensive for large datasets. In addition, K-Nearest Neighbors (KNN) regression was included ^56^. It relies on the proximity of data points in feature space to make predictions. KNN is effective in capturing local structures but may struggle with global generalization. Several linear and regularization-based models were also evaluated, including Bayesian Ridge Regression and ElasticNet Regression^51,52^. Bayesian Ridge is a probabilistic linear model that introduces regularization through a Bayesian prior, making it more robust to collinearity in the dataset. ElasticNet combines L1 (Lasso) and L2 (Ridge) penalties, balancing input variable selection with coefficient shrinkage to prevent overfitting. Decision Tree Regression ^50^ was included as a baseline tree-based model, providing a simple interpretable alternative to ensemble methods. Although useful for understanding decision boundaries, single decision trees often overfit the data and lack the predictive power of ensemble methods.

#### Training

Each AI/ML model was trained on the dataset using 5-fold cross-validation. The models were first trained on four subsets of the data and then validated on the remaining subset, with the process repeating five times to obtain an average performance score ^57^. R² Score (Coefficient of Determination) was calculated to measure the proportion of variance explained by the model ^58^. R² score close to or like 1 indicates a better fit. Mean Absolute Error (MAE) measures the average absolute difference between predicted and actual values ^59^. Lower values indicate better accuracy. Mean Squared Error (MSE), not shown in graph, was visible in console when running script. MSE is similar to MAE but penalizes larger errors more heavily^59^. Hyperparameters for tree-based models were optimized to balance bias and variance. For example, Random Forest and Extra Trees were trained with 300 estimators (trees) and a depth limit of 6 to prevent overfitting. Gradient Boosting and CatBoost were fine-tuned with a learning rate of 0.05–0.1 to capture smooth nonlinear patterns. Gaussian Process Regression was configured with a Radial Basis Function (RBF) kernel for smooth nonlinear patterns.

#### Analysis

The analysis was conducted in the Visual Studio Code software was used with Python-based computational tools for data processing, machine learning modeling, and visualization. Data handling and preprocessing were performed using Pandas for tabular data manipulation and NumPy for numerical computations ^60,61^. For predictive modeling, various machine learning algorithms were applied using Scikit-Learn ^62^ and CatBoost ^48^. Matplotlib was used to visualize the fitted models against the original dataset ^63^. The predictions generated by each model were visualized against the actual data points to assess their alignment with the observed trend.

## Data availability

Additional data in connection with the figures are available from the corresponding author on reasonable request.

## Acknowledgements

This study was supported by grants from the Faculty of Medicine and Health Sciences and Samarbeidsorganet (The cooperation body between Helse-Midt Norge RHF and universities and colleges in the region) and P30CA013696 (AKR, TCW) and its Proteomics and Structural Biology Shared Resource. We would like to thank Drs. David Tuveson and Park Young at Cold Spring Harbor Laboratory, New York for offering scientific advice and murine PDAC organoids; Drs. Xin Cai and Wei Wang at NTNU and Linda Trobe Dorg at UiO for scientific and technical advice/assistance particularly on cultures of cells and organoids and human PDAC materials.

## Author contributions

M. R. and E.P.G. conceived the presented data. M.R. E.P.G., H-L.R., A.S, L.H., L.H., N.T.S., A.A., M.K.S, M.A., C.S.V., S.K.B. A.R. contributed to model preparations and the data analysis. C-M.Z., G.Q. and T.C.W. supervised this PhD project by M.R. D.C. supervised this MSc project by E.P.G. C-M.Z. supervised this MSc by A.A. C-M.Z. and D.C. conceived this study and helped M.R. to write the manuscript and edited the manuscript with input from all co-authors.

## Competing interests

The authors declare no competing interests. Of note, part of this study was included in a MSc thesis entitled “Organoids as a potential model for studying pancreatic ductal adenocarcinoma” by Anne Aarvik (https://ntnuopen.ntnu.no/ntnu-xmlui/handle/11250/3135404, in which full text not available for open access).

## Notes

### Competing Interest Statement

The authors have declared no competing interest.

